# Synaptic plasticity onto inhibitory neurons as a mechanism for ocular dominance plasticity

**DOI:** 10.1101/280511

**Authors:** Jacopo Bono, Claudia Clopath

## Abstract

Ocular dominance plasticity is a well-documented phenomenon allowing us to study properties of cortical maturation. Understanding this maturation might be an important step towards unravelling how cortical circuits function. However, it is still not fully understood which mechanisms are responsible for the opening and closing of the critical period for ocular dominance and how changes in cortical responsiveness arise after visual deprivation. In this article, we present a theory of ocular dominance plasticity. Following recent experimental work, we propose a framework where a reduction in inhibition is necessary for ocular dominance plasticity in both juvenile and adult animals. In this framework, two ingredients are crucial to observe ocular dominance shifts: a sufficient level of inhibition as well as excitatory-to-inhibitory synaptic plasticity. In our model, the former is responsible for the opening of the critical period, while the latter limits the plasticity in adult animals. Finally, we also provide a possible explanation for the variability in ocular dominance shifts observed in individual neurons and for the counter-intuitive shifts towards the closed eye.

## INTRODUCTION

Throughout development, sensory cortex can experience periods of heightened sensitivity to sensory inputs. The rewiring of neuronal networks is very flexible during these periods, but there is less such plasticity otherwise. Having normal sensory experiences during these periods is crucial for a healthy maturation of the brain and they are therefore called critical periods (CP).

A well studied example is the critical period for ocular dominance (OD) in primary visual cortex (V1). In the visual pathway, inputs from both eyes usually converge onto the same neuron for the first time in V1, although a fraction of thalamic neurons already exhibits binocularity in mice [Jaepel et al., 2017, Sommeijer et al., 2017, Jeon and Kuhlman, 2017]. The extent to which a neuron’s visually-evoked activity is dominated by one of the eyes is called ocular dominance (OD) and is often quantified by the ocular dominance index (ODI). In each hemisphere of mice V1, the overall response to the contralateral eye is roughly twice as high as that to the ipsilateral eye, but individual neurons display a broad range of ODI values.

During a limited period early in the development, closure of one eye for multiple days triggers a shift in neuronal responses towards the open eye. In mice, this critical period spans about ten days, starting around postnatal day 20. The changes in neuronal responses following this monocular deprivation (MD) can be roughly separated into two phases. In a first phase, observed during the first three days of deprivation, the responses to the closed eye are depressed while responses to open-eye inputs remain similar. This effect is often called response depression. For longer deprivations, a second phase follows where the neuronal responses to the open eye are increased, called response potentiation. In this second phase, the neuronal activity caused by the closed eye also increases, but to a lesser extent [Frenkel and Bear, 2004]. Further insights into the working of ocular dominance plasticity are uncovered by studying other deprivation paradigms. Firstly, binocular deprivation (BD) does not lead to OD shifts, hinting at some level of competition depending on the strength or coherence of the inputs from both eyes. Secondly, monocular inactivation (MI) abolishes the rapid response depression, suggesting that this response depression is activity dependent, relying on spontaneous activity and residual activity caused by light travelling through the closed eyelid during MD [Frenkel and Bear, 2004].

Before and after the critical period, the effects of monocular deprivation on ocular dominance plasticity are either reduced or not observed at all. In pre-CP mice, monocular deprivation leads to a decrease in activity from both eyes, thus not changing their relative strengths and not affecting the overall ocular dominance [Smith and Trachtenberg, 2007]. In adult mice, the response depression after short monocular deprivation is not observed. However, longer deprivation still leads to the response potentiation of the open eye, and hence a certain shift in ocular dominance can still be observed [Sawtell et al., 2003].

A key player in regulating the opening of the critical period is the maturation of inhibition. GAD95-KO mice, which exhibit an impaired *γ*-aminobutyric acid (GABA) release, never experience a critical period and visual cortex remains in a juvenile state. A critical period can be opened in these mice once per lifetime after diazepam infusion, which restore the GABA release. Similarly, diazepam infusion before normal CP onset can accelerate the start of the CP in wildtype mice [Fagiolini and Hensch, 2000]. Furthermore, a recent experimental study investigated changes in cortical layer II/III excitation and inhibition in juvenile animals after only 24 hours of deprivation [Kuhlman et al., 2013]. The authors found that the firing rate of parvalbumin-positive (PV+) inhibitory neurons is decreased at that time, while the firing rate of excitatory neurons is increased. Moreover, they show that this decreased inhibition is predominantly mediated by a reduction in excitatory drive from layer IV and V to these PV+ neurons. Interestingly, this reduction of inhibition is not observed in adult animals. The authors then linked this effect to the OD shift, by showing how pharmacological enhancement of inhibition during the critical period prevents any OD plasticity, while pharmacological reduction of inhibition in the adult animals results in an OD shift towards the open eye. It was therefore postulated that OD plasticity depends on the increased firing rate of open eye inputs, caused by a transient reduction in inhibition.

The mechanisms behind the closure of the critical period remain more enigmatic. However, several manipulations can reopen a window for OD plasticity in adult mice. Firstly, reducing inhibition was shown to enhance the OD plasticity caused by monocular deprivation [Harauzov et al., 2010, Kuhlman et al., 2013]. Related to this, adult mice in enriched environments were shown to have reduced levels of inhibition and OD plasticity [Greifzu et al., 2014]. Finally, high contrast stimulation during deprivation also leads to OD shifts [Matthies et al., 2013], suggesting that enhancing visually evoked responses of the open eye could be functionally similar to reducing cortical inhibition. Other mechanisms that have been implied with the ending of the critical period are changes in the extracellular matrix [Pizzorusso, 2002] and the pruning of silent synapses [Huang et al., 2015]. Taken together, experimental results hint at a visual-experience dependent maturation of V1, where normal visual stimuli are necessary to shape the network connectivity.

In this article, we propose a model for the first phase after deprivation, coinciding with the response depression phase under MD. We follow the hypothesis that a reduced inhibition is the key to allow for plasticity. More specifically, we model a neuronal network and propose synaptic plasticity principles that are able to reproduce many of the phenomena discussed above. In our model, excitatory-to-inhibitory plasticity is responsible for a rapid reduction in inhibition during the CP, which in turn enables a shift in ocular dominance. Our model is consistent with experimental results observed under MD of the contra- and ipsilateral eyes, under MI and under BD. Furthermore, we discuss possible mechanisms underlying the opening and closing of the critical period, and reinstatement of plasticity. Finally, our model provides a possible explanation to why some neurons shift counter-intuitively towards the closed eye and why these neurons tend to have lower firing rates.

## RESULTS

### 1. Hypothesis for unifying juvenile and adult OD plasticity

The experimental observation that increasing inhibition in pre-CP animals [Fagiolini and Hensch, 2000] and decreasing inhibition in adult animals [Harauzov et al., 2010] allows for OD plasticity, naturally leads to a two-level inhibition hypothesis. More specifically, a first increase in inhibition would open the critical period and a further increase of inhibition would close it. However, the results by Kuhlman et al. [Kuhlman et al., 2013] allow for a different interpretation. Indeed, the authors showed that even during the critical period, a reduction of inhibition is necessary to observe OD plasticity. The authors therefore proposed that the increased levels of inhibition are crucial in opening the critical period because a subsequent reduction of inhibition can amplify the open-eye excitatory activity. Moreover, stimulating adult animals with high contrast gratings also leads to fast OD plasticity [Matthies et al., 2013]. We can then unify all the results by stating that an OD shift towards the open eye is possible when the open-eye responses are transiently increased. This could be either by reducing inhibition, or by enhancing excitation. Thus, we hypothesise the following:

- The loss of input after monocular deprivation shifts both open-eye and closed-eye pathways towards depression. No OD shift would be observed at this point.
- Boosting the open-eye inputs, for example by a quick reduction inhibition, pushes the open-eye, but not the closed-eye inputs over a threshold for potentiation. This leads to the depression of closed-eye inputs while maintaining open eye responses.

To test this hypothesis in simulated networks, we first consider a simplified model where a single neuron representing a layer II/III pyramidal cell receives feedforward excitatory input from a population of layer IV neurons and feedforward inhibition from one inhibitory neuron (Fig. 1a). The feedforward excitatory synapses onto the layer II/III neuron are plastic, while other synapses are static. Assuming *ρ*_pre_ and *ρ*_post_ are the presynaptic and postsynaptic firing rates respectively, we use the following Hebbian excitatory learning rule (see Methods). If the the product (*ρ*_pre_ · *ρ*_post_) caused by an input exceeds a threshold *θ_H_*, the synaptic weight is increased by an amount *η*. If this value remains below *θ_H_*, the value is decreased by *η*. The parameter *θ*_*H*_ therefore is a constant threshold separating synaptic depression from synaptic potentiation, and *η* is the learning rate. In this way, layer IV synapses with various ODIs onto the same layer II/III neuron will lead to different values for (*ρ*_pre_ · *ρ*_post_) after deprivation. Indeed, a layer IV neuron that is dominated by the deprived eye, will be left with a small value for *ρ*_pre_ after MD, while a layer IV neuron dominated by the open eye will be relatively unaffected and have a high *ρ*_pre_ (Fig. 1b). Moreover, MD will also reduce the postsynaptic firing rate *ρ*_post_ of the layer II/III neuron by an amount depending on its ocular dominance index. Finally we initialize the feedforward excitatory weights at the upper bound.

**Figure 1.**
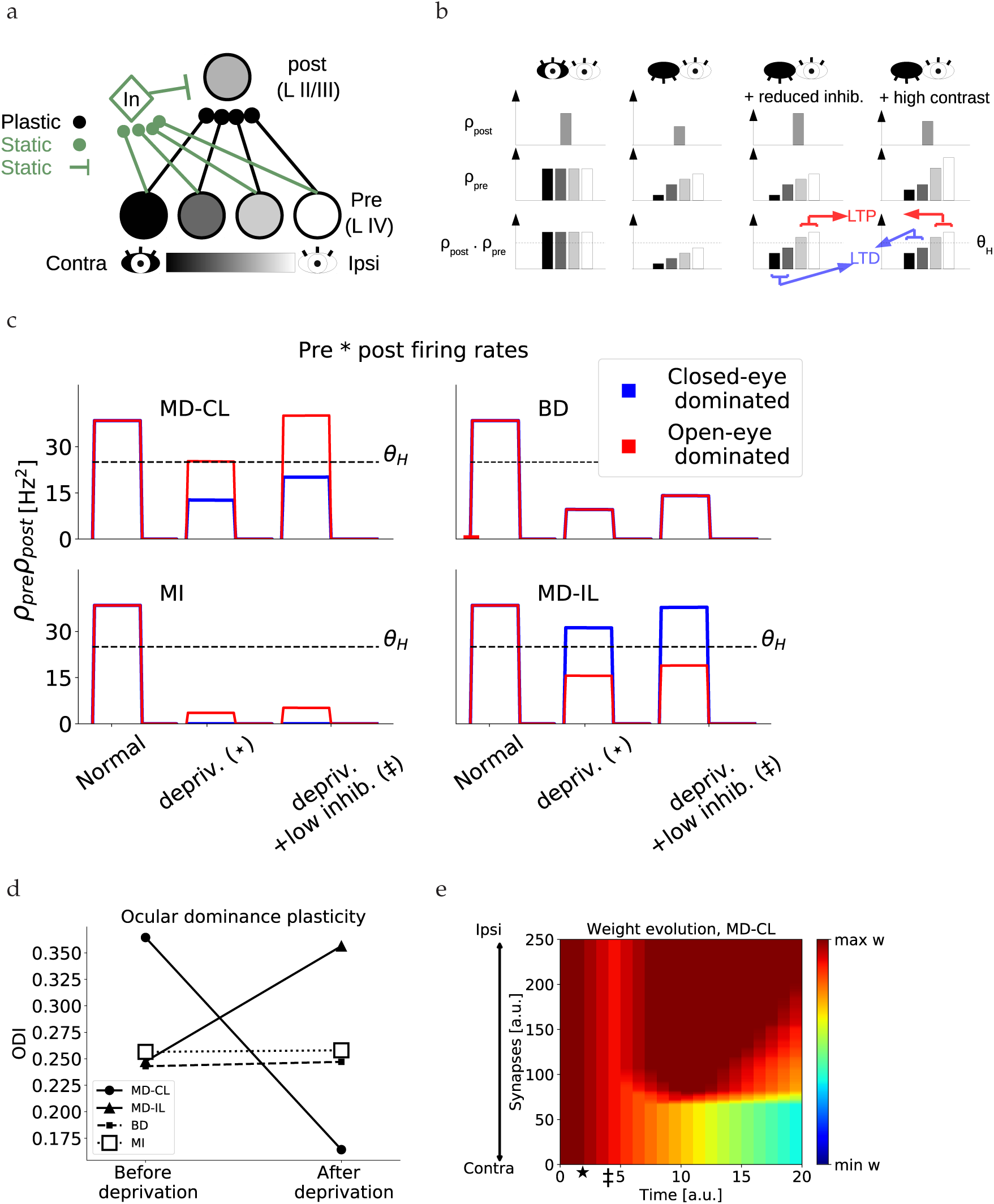
Single-threshold model. (a) Schematic of the model. One postsynaptic neuron receives inputs from a population of presynaptic neurons, each with different ODI. We use a grey-scale color scheme where darker colors denote contralateral-eye dominated neurons and brighter colors denote ipsilateral-eye dominated neurons. (b) Schematic of preand postsynaptic firing rates under normal rearing (left), monocular deprivation of the contralateral eye (second from left), monocular deprivation with a reduced inhibition (second from right) and monocular deprivation with high contrast inputs (right). The bottom row depicts the product between preand postsynaptic firing rates, and the threshold *θ* separating LTD from LTP. (c) Example traces of the product of presynaptic and postsynaptic firing rates of the neurons during monocular deprivation of the contralateral eye (MD-CL), binocular deprivation (BD), monocular inactivation of the contralateral eye (MI) and monocular deprivation of t5he ipsilateral eye (MD-IL). For red and blue traces, the product is taken with the most open-eye dominated and closed-eye dominated presynaptic neuron respectively.(d) Ocular dominance index of the postsynaptic neuron at the beginning of the simulation versus the end. (e) Evolution of synaptic weights over time, in the case of monocular deprivation of the contralateral eye. The star denotes the onset of deprivation, the double dagger denotes the onset of reduced inhibition. Corresponding neuronal activations are shown in panel c.

In this simplified model, we first assume that monocular deprivation pushes all possible (*ρ*_pre_ · *ρ*_post_) values into the long-term depression (LTD) regime by reducing *ρ*_pre_ and *ρ*_post_. Since all synapses are depressed equally, this does not affect the relative response strength between the eyes and therefore leaves the ODI unaltered. Secondly, a reduction of inhibition can rescue the original postsynaptic firing rate and hence shifts only the (*ρ*_pre, open_ · *ρ*_post_) above the long-term potentiation (LTP) threshold *θ_H_*. Here, by *ρ*_pre, open_ we mean the presynaptic rates corresponding to open-eye dominated neurons. Only after this reduction of inhibition, the ocular dominance shifts by depressing the closed-eye inputs while maintaining the open-eye inputs (Fig. 1b,d).

### 2. Simulating different types of deprivation

To simulate this simplified model, we connect 250 presynaptic neurons to one postsynaptic excitatory neuron and one inhibitory neuron, each modelled as rate units. The presynaptic neurons have a broad range of ODIs (see Methods and Supp. Fig. 6a). In this simplified model, only the layer IV to layer II/III excitatory inputs are plastic and initialized at the upper bound. The layer IV neurons are activated by visual inputs from both eyes and a background input (see Methods). When both eyes are open, all these excitatory inputs are in the LTP regime and therefore remain at the upper bound. We then simulate monocular deprivation of the contralateral eye (MD-CL) by reducing the input of the contralateral eye to zero. The closure of the eye therefore reduces the firing rates and all synapses undergo LTD (Fig. 1c, top left). We subsequently reduce the feedforward excitatory-to-inhibitory connections to a third of the initial value. This reduction of inhibition leads to a recovery of the original postsynaptic excitatory firing rate, consistent with the data from Kuhlman et al. [Kuhlman et al., 2013]. Now only the feedforward connections from presynaptic neurons dominated by the closed eye are depressed, while the potentiation of the open eye pathway brings the respective synapses back to the upper bound (Fig. 1c, top left). This depression of the closed-eye pathway ultimately leads to an OD shift toward the open eye (Fig. 1d,e).

The same model can also reproduce the lack of OD plasticity after binocular deprivation. In this case, both eyes are sutured and therefore all inputs are reduced to a third of the original values. Since all presynaptic firing rates are attenuated by an equal amount, all (*ρ*_pre_ · *ρ*_post_) have the same value. This ensures that the open-eye and closed-eye inputs will always have the same direction of plasticity. In our case, they are all in the LTD region and therefore depressed (Fig. 1c top right, d).

In the case of monocular inactivation of the contralateral eye, TTX injection in the retina abolishes all neuronal activity. This is in contrast with MD-CL, where spontaneous activity is present and some light can travel through the sutured eyelid. Experimentally, no ocular dominance shift is observed after monocular inactivation [Frenkel and Bear, 2004], suggesting that the residual activity is important. Similar to BD, the total amount of neuronal activity is lower under MI than under MD-CL. However, unlike BD the presynaptic inputs strengths are now variable, depending on the ODI of the respective input. This resembles the situation under MD-CL, but with all input strengths shifted to lower values. With an appropriate choice for the threshold *θ_H_*, we can therefore still obtain that all synapses are depressed, and hence no OD shift is observed in our postsynaptic neuron (Fig. 1c bottom left, d).

Finally, in the case of monocular deprivation of the ipsilateral eye (MD-IL), we follow the same reasoning as in the MD-CL case. Closed-eye inputs fall below *θ_H_* and open-eye inputs remain above, leading to a shift towards the contralateral eye (Fig. 1c bottom right, d). This shift is in agreement with [Sato and Stryker, 2008, Mrsic-Flogel et al., 2007]

### 3. Heterogeneity in OD shifts

Recent work by Rose et al. [Rose et al., 2016] uncovered a substantial degree of heterogeneity in OD plasticity of individual neurons after monocular deprivation. About 40% of the neurons in layer II/III do not show any particular plasticity, while the amount and direction of the shift in the remaining 60% is variable. Indeed, some neurons even shift their responses counter-intuitively towards the closed eye. The latter neurons were shown to have lower visually-evoked activities and the counter-intuitive shift was caused by a depression of the open-eye inputs. Counterintuitive shifts were also observed in a study with cats, where a global counter-intuitive shift towards the closed eye occurred after increasing the inhibition during the monocular deprivation [Reiter and Stryker, 1988].

Most neurons in layer IV receive inputs coming from both eyes and project to layer II/III neurons. If these layer IV neurons all have very similar ocular dominance, all synapses will be modified in a similar way in our simplified model and no OD shift will be observed. Indeed, it is highly unlikely that the threshold *θ_H_* will fall somewhere within this narrow distribution (Fig. 2a). For layer II/III neurons to show OD plasticity, the difference between the minimum and maximum ODI of incoming layer IV inputs must therefore be large enough (Supp. Fig. 2a,b). Assuming a variety of OD distributions for the inputs would therefore suffice to reproduce both non-plastic neurons and neurons shifting towards the open eye with different magnitude, but not the counterintuitive shifts towards the closed eye.

**Figure 2.**
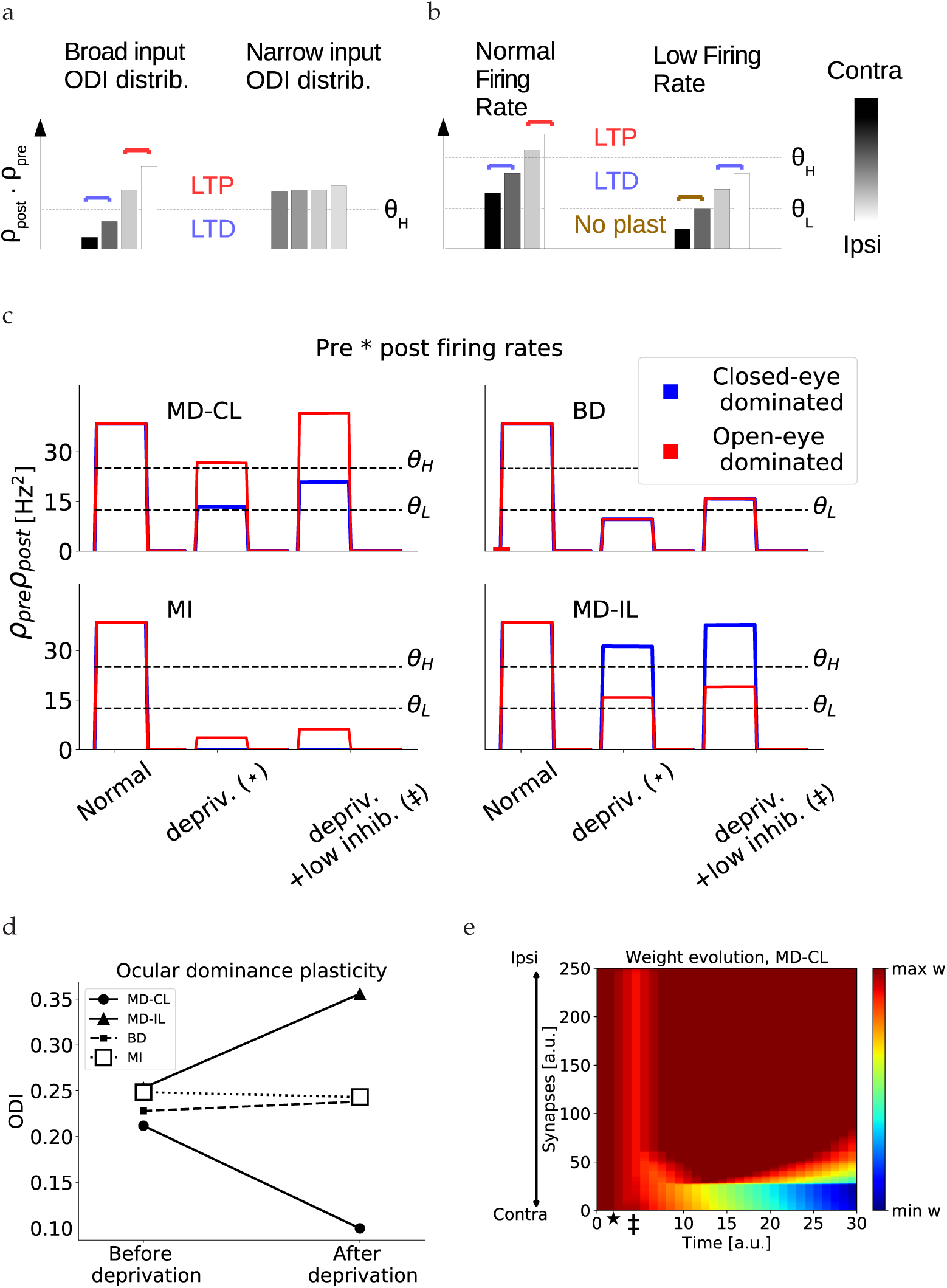
Double-threshold model. (a) Schematic of presynaptic multiplied by postsynaptic firing rates after monocular deprivation, and assuming different ODI distributions of input neurons. If the ODI distribution is too narrow, it is harder for the threshold *θ* to separate more closed-eye dominated inputs from more open-eye dominated inputs. (b) Schematic of presynaptic multiplied by postsynaptic firing rates after monocular deprivation, and assuming heterogeneous postsynaptic firing. Postsynaptic neurons that receive weaker inputs and fire at lower rates, lead to a depression of open-eye dominated inputs while leaving closed-eye inputs unaltered. These neurons show a counter-intuitive shift towards the closed eye. (c) Examples of the product of presynaptic and postsynaptic firing rates of the neurons during MD, BD and MI. For red and blue traces, the product is taken with the most open-eye dominated and closed-eye dominated presynaptic neuron respectively. (d) Ocular dominance index of the postsynaptic neuron at the beginning of the simulation versus the end. (e) Evolution of synaptic weights over time, in the case of monocular deprivation of the contralateral eye. The star denotes the onset of deprivation, the double dagger denotes the onset of reduced inhibition. Corresponding neuronal activations are shown in panel c.

In order to reproduce these counter-intuitive shifters, we adapt our plasticity rule to contain a second threshold. Besides the threshold separating the LTD region from the LTP region, we introduce a lower threshold which separates a no-plasticity region from the LTD region. We can then understand the counter-intuitive shifters as follows. Neurons receiving low-rate input and/or firing at low rates exhibit smaller values of (*ρ*_pre_ · *ρ*_post_) compared to Fig. 1. Subsequently, the closed-eye pathway could fall below the lower threshold for plasticity and would not be altered, while the open-eye pathway would be in the depression regime (Fig. 2b). This results in a counter-intuitive shift where the closed eye gains strength relative to the open eye.

Finally, with this modified learning rule, we set our thresholds so that the (*ρ*_pre_ · *ρ*_post_) fall below the threshold for plasticity immediately after deprivation but before the reduction of inhibition. With this choice, the synapses are unaltered immediately after deprivation, as opposed to all being depressed as in Fig. 1b. Since both scenarios would not change the relative strength of the two eyes until the inhibition is reduced, they both agree with experiments. In practice, because of the variety of ODIs in the inputs, some inputs dominated by the open eye may fall in the depression regime (Fig. 2c, top left). Since we will assume that excitatory-to-inhibitory plasticity has a faster action then excitatory-to-excitatory plasticity, this short period of open-eye LTD has a negligible and transient effect Fig. 2e.

### 4. Larger network simulations

The simplified models discussed in the previous sections only included feedforward excitatoryto-excitatory (E-to-E) plasticity onto a single layer II/III neuron. We now expand this framework to a population of layer II/III neurons, while adding excitatory and inhibitory plasticity in all connections (Fig. 3a). The E-to-E plasticity rule remains the same as before, with a low threshold *θ_L_* below which no plasticity occurs, and a high threshold *θ_H_* separating synaptic depression from potentiation (Fig. 3b). For the E-to-I plasticity rule, we require that it is not too selective and that synaptic depression is induced after monocular deprivation. The first requirement follows from the experimental observation that inhibitory neurons are broadly tuned for orientations [Hofer et al., 2011], while the second requirement is necessary for ocular dominance plasticity in our model. We therefore choose to model E-to-I plasticity using a modified version of the BCM-rule [Bienenstock et al., 1982] with hard upper bounds on the synaptic weights. We choose the target firing rate to be dependent on the visual experience. For normal binocular vision, a high target firing rate enables strengthening of inhibition and a reduced selectivity. For all forms of deprivation, we reduce the target firing rate to 10% of its original value, ensuring a depression of all E-to-I connections. In the cases of monocular deprivation, this depression is followed by a recovery of the open-eye inputs (Fig. 4c). Raising animals in complete darkness would therefore never lead to a maturation of inhibition, as observed experimentally [Benevento et al., 1992, Fagiolini et al., 1994]. Finally, the I-to-E plasticity rule is a modified version of the rule proposed in Vogels et al. [Vogels et al., 2011], ensuring each excitatory neuron does not exceed a maximal firing rate (see details in the Methods section).

**Figure 3.**
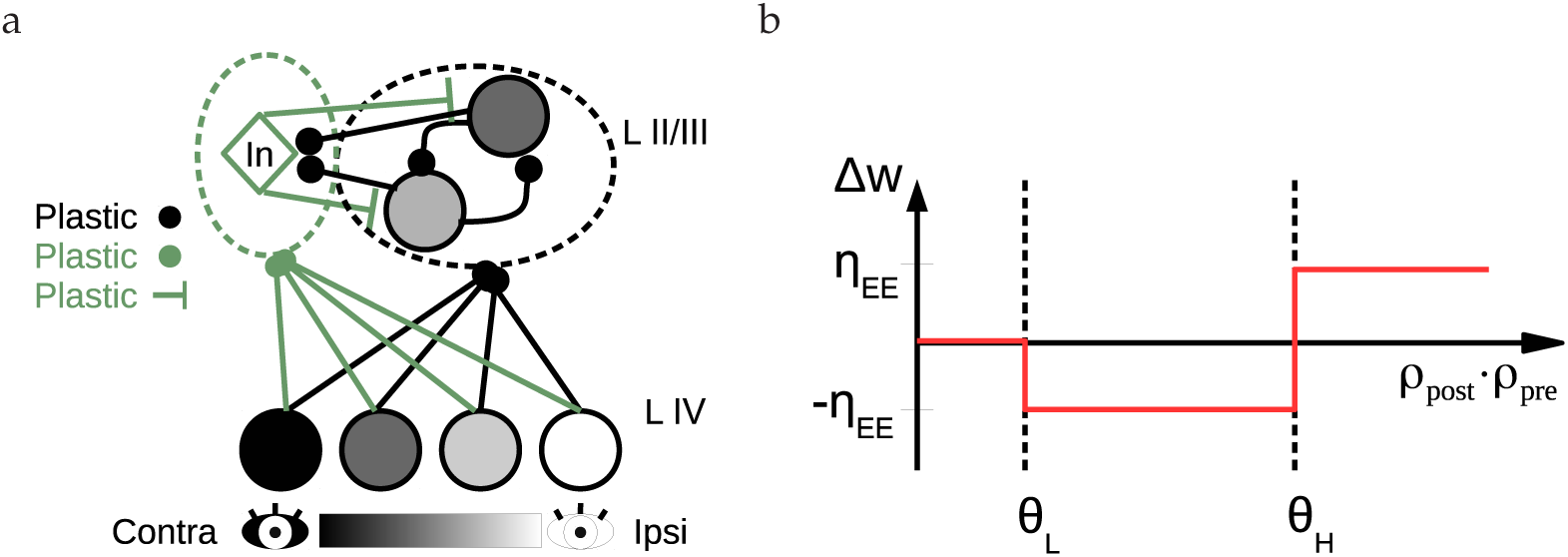
Schematic of the network and excitatory learning rule. (a) A cartoon of the network architecture. A population of presynaptic (layer IV) neurons with various ODIs makes feedforward connections onto layer II/III excitatory and inhibitory neurons. Within the layer II/III, excitatory neurons have recurrent connections both to other excitatory neurons as well as to the inhibitory neurons, which in turn project back onto the excitatory neurons. (b) Schematic of the E-to-E plasticity rule. Low values for the product of presynaptic and postsynaptic rates do not lead to plasticity. Intermediate values result in synaptic depression, and high values in synaptic potentiation.

**Figure 4.**
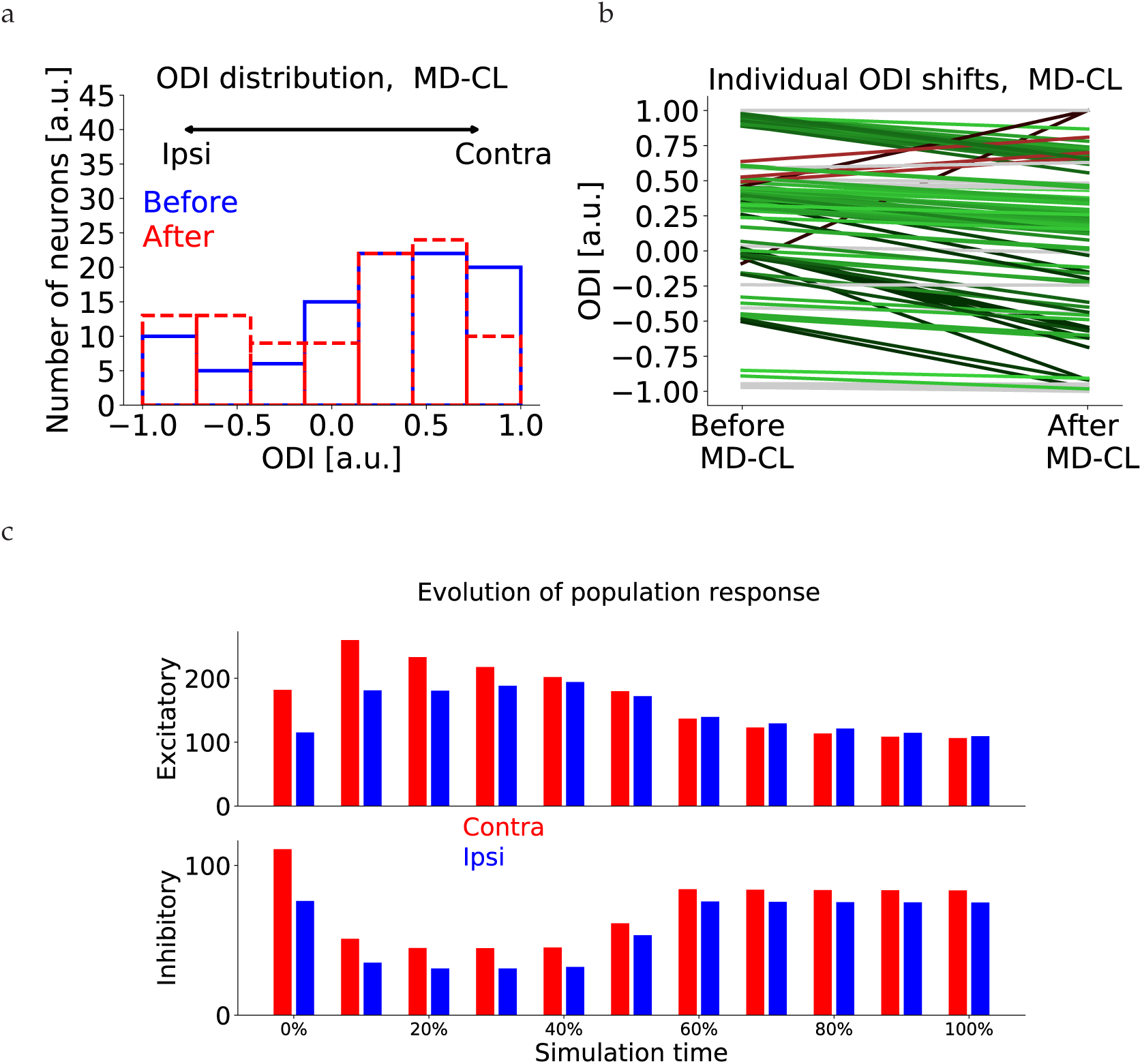
Network simulations of MD. (a) Distribution of ODI values of layer II/III neurons before MD (blue) and after MD (red). (b) Individual OD shifts for all layer II/III neurons after MD. Green lines denote shifts towards the open eye. Red lines denote counter-intuitive shifts towards the closed eye. (c) Excitatory and inhibitory population response to contraand ipsilateral eye over time. The 0% denotes the start of deprivation. A quick reduction of inhibition is followed by a recovery of mainly the open-eye inhibition, as observed in [Kuhlman et al., 2013].

We assume a variety of ocular dominances in both layer IV and layer II/III neurons (see Methods and Supp.Fig. 6a), and a variety of excitatory firing rates. Moreover, we divide the layer IV neurons into five groups that are activated separately. These groups mimic the encoding of different input features, for example differently oriented lines within a receptive field. After an initial phase of the simulation where excitatory connections reach either upper or lower bounds, we simulate MD-CL, MD-IL, BD and MI. Similar to the simplified model, we simulate MD by abolishing the visual input from the corresponding eye while maintaining the background input. At the end of the simulation, the mean response of layer II/III neurons shifted towards the open eye (Fig. 4a,b, Fig. 5c). This shift is mediated by a depression of closed-eye inputs, while openeye inputs remain roughly the same (Fig. 4b,c). Individual neurons show a variety of OD shifts, and some neurons shift counter-intuitively towards the closed eye (Fig. 4b). When plotting the individual shifts versus the firing rate, it is clear that the counter-intuitive shifters are neurons with lower-than-average firing rates (Fig. 5a). We then simulate MI by completely abolishing both the contralateral input and background input, and BD by abolishing contra- and ipsilateral inputs but not the background. Since the firing rates of all neurons are now substantially lower, no OD shift is observed (Fig. 5c, Supp. Fig. 4). Finally, in the cases of MD, we observe that inputs driven exclusively by the closed eye do not show any OD shift (Fig. 5d). Indeed, for these neurons the firing rates are more significantly reduced and similar to the case of BD. This dependence of OD shift on initial ODI was experimentally observed in [Mrsic-Flogel et al., 2007].

**Figure 5.**
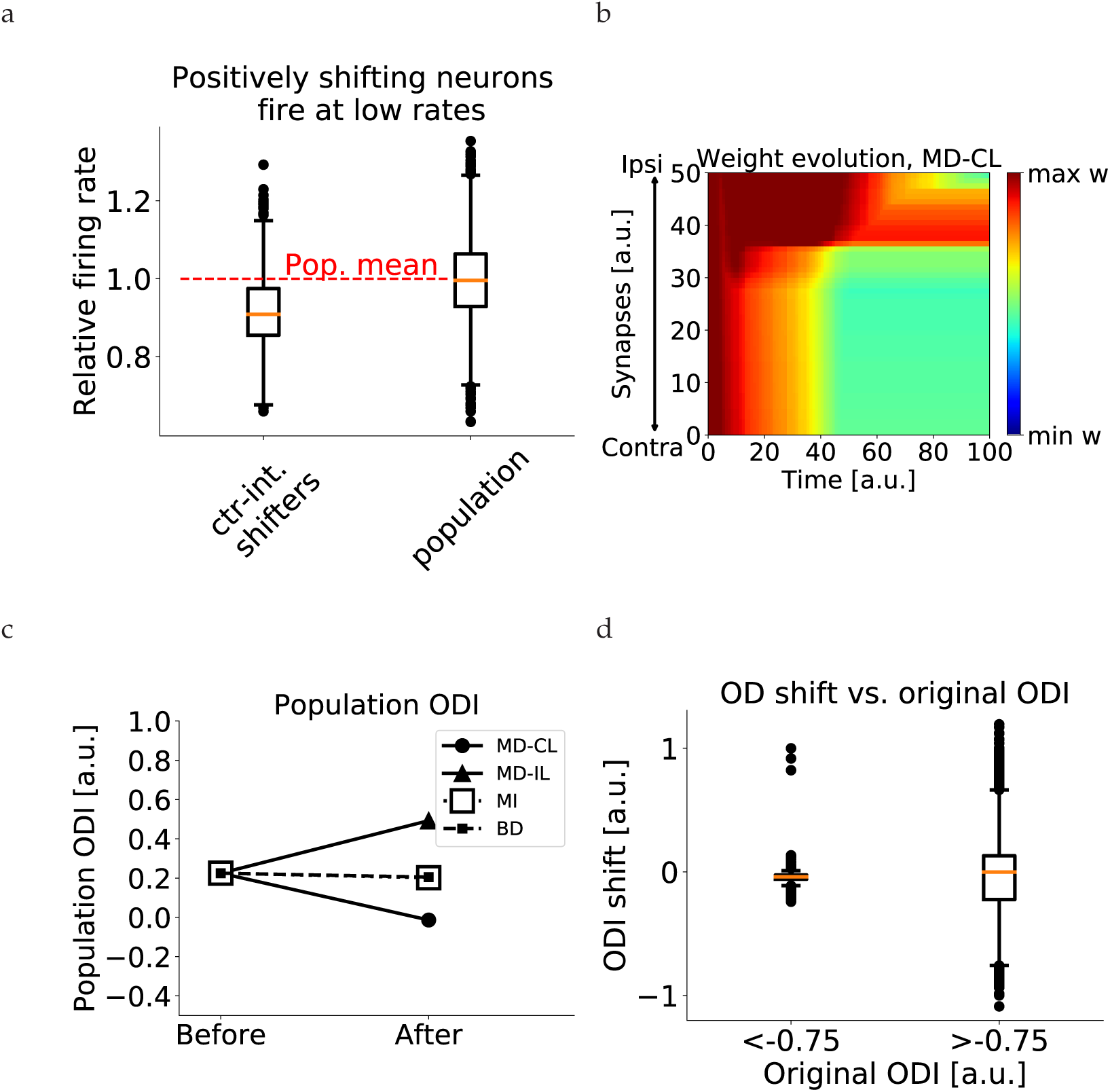
Network simulations of deprivation. (a) Relative firing rate distribution (normalized to population mean) for counter-intuitive shifters and for the total population. We consider a neuron to be a counter-inuitive shifter if the ODI difference is larger than 0.15. The population consists of 5000 neurons (50 network simulations of MD-CL with 100 neurons each). For the counter-intuitive shifters, most of the firing rates lie below the mean. (b) Evolution of synaptic weights over time for one neuron in the network, in the case of monocular deprivation of the contralateral eye. Similarly, for MD-IL, BD and MI, see supp. Fig. 4. (c) Population ODI shifts for all forms of deprivation. Under both types of MD, the population shifts towards the open eye. No shift is observed after BD and MI. (d) 5000 neurons (50 network simulations of 100 neurons) are divided according to original ODI. The boxplots show the distribution of OD shifts for both groups. Ipsilaterally dominated neurons show no response depression after MD-CL.

### 5. Onset and ending of the critical period

In order to simulate the maturation of the network, we firstly assume that it starts from an ’immature’, pre-critical-period state. We assume that the inhibitory and excitatory recurrent connections are still weak and the E-to-I connections start close to the minimum bound. In agreement with experimental data from Hofer et al. [Hofer et al., 2011], the choice of our learning rules ensure that E-to-E connections are input-selective while E-to-I connections are unspecific. Moreover, during the development, excitatory neurons increase their input selectivity over time while inhibitory neurons broaden their input tuning (Fig. 6c). This can be understood as follows. Both inhibitory and excitatory neurons start with a small bias for one input and therefore have a low selectivity index. However, our selective E-to-E rule ensures that excitatory neurons only develop strong connections with similarly tuned neurons (Fig. 6d, Supp. Fig. 3a), while the unselective E-to-I rule ensures that inhibitory neurons strengthen all incoming connections (Fig. 6e, Supp. Fig. 3b). This evolution is in qualitative agreement with experimental observations of input selectivity in juvenile mice [Kuhlman et al., 2011].

**Figure 6.**
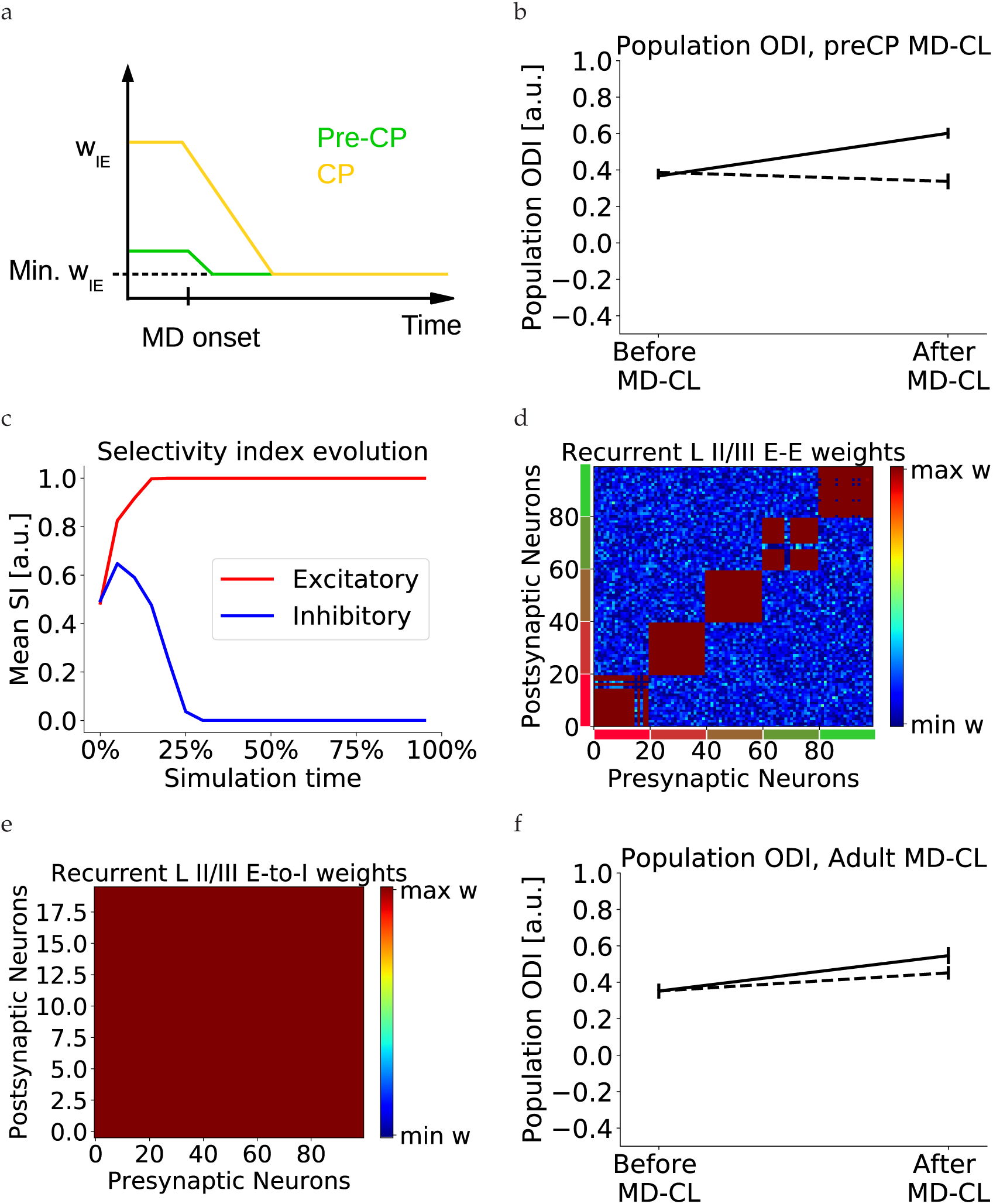
Pre-CP and ending of the CP. (a) In the pre-CP phase, the inhibition is still weak and close to the minimum value. Unlike CP mice, the inhibition cannot be sufficiently reduced before CP onset. (b) MD-CL does not lead to an OD shift towards the open eye in pre-CP mice. An OD shift towards the closed eye is observed (solid line), which is reduced when assuming a lower visual-to-background activity ratio (dashed line). (c) In this pre-CP phase, the input selectivity index of excitatory neurons increases, while inhibitory neurons broaden their selectivity. This is in qualitative agreement with experimental data [Kuhlman et al., 2011]. (d) Neurons are grouped according to input feature preference (different colors next to axes denote different input preference). Recurrent E-to-E weights are specific: only synapses from neurons with similar input preference are strong, while other recurrent inputs are weak. (e) Recurrent E-to-I weights are unspecific, synapses from all input groups are at the maximum bound. (f) We simulate adult networks by preventing any E-to-I plasticity. MD-CL does not lead to an OD shift towards the open eye. An OD shift towards the closed eye is observed (so1l4id line), which is reduced when assuming a higher visual-to-background activity ratio (dashed line).

We can then simulate both the pre-CP and adult networks by assuming that the inhibition is unable to amplify the open-eye activity after deprivation. For the pre-CP case, the inhibitory activity cannot be sufficiently reduced after monocular deprivation because of the immature levels of inhibition in the pre-CP period (Fig. 6a). For the adult case, one candidate mechanism could be the consolidation of these E-to-I synapses blocking E-to-I plasticity. When simulating both cases, no OD shift towards the open eye is observed. In fact, since some inputs dominated by the open eye fall in the depression regime, we notice a slight global counter-intuitive shift (Fig. 6b,f solid lines). This shift is eliminated by assuming that the background-to-visual ratio is higher in the pre-CP and lower in the adult, as observed in [Toyoizumi et al., 2013] (Fig. 6b,f dashed lines).

## Discussion

In this article, we simulated a simplified model of a layer II/III network in primary visual cortex. Our model is able to reproduce several experimentally observed features of the critical period for ocular dominance. In particular, we simulated changes caused by monocular deprivation, binocular deprivation and monocular inactivation. Furthermore, we discuss possible mechanisms for the onset and the end of the critical period. Our model therefore provides possible mechanistic insights into the development of cortical areas and the associated learning rules, which could be tested experimentally.

Our aim was to acount for the effects of short deprivation (up to 3 days) on the ocular dominance of layer II/III neurons. We did not include longer deprivations in our model (more than 3 days), when response potentiation is observed. In this case, responses to both eyes but mainly to the open eye start to increase. Therefore homeostatic plasticity mechanisms are a likely candidate to explain this second phase of OD plasticity [Turrigiano et al., 1998, Mrsic-Flogel et al., 2007, Espinosa and Stryker, 2012]. Furthermore, the study observing counter-intuitively shifting cells [Rose et al., 2016] was performed on adult mice. However, these mice were kept in enriched environments and stimulated with high contrast inputs, both known to enable a juvenile-like plasticity [Greifzu et al., 2014, Matthies et al., 2013].

We only considered plasticity in connections from layer IV to layer II/III and within layer II/III. Therefore, we did not take into account experimentally observed OD shifts in the thalamic relay neurons [Sommeijer et al., 2017, Jaepel et al., 2017] and layer IV neurons [Gordon and Stryker, 1996]. Since these areas are upstream of layer II/III, a naive explanation could be that the shift in layer II/III is fully accounted for by the shift in the inputs to this layer. However, Gordon and Stryker [Gordon and Stryker, 1996] described how a larger OD shift is observed in layer II/III compared to layer IV neurons, and similarly a larger shift in layer V/VI is observed compared to layer II/III. Considering the canonical flow of sensory inputs, from thalamus to layer IV, further to layer II/III and finally to layers V/VI, this result suggests that plastic changes happen at each stage and accumulate over layers.

An increased inhibition is necessary in our model to open the critical period. This is because weak inhibition cannot be reduced sufficiently to rescue excitatory firing rates after monocular deprivation. Our hypothesis differs from previous theories on the opening of the critical period, which did not take into account the transient reduction of inhibition observed by Kuhlman et al. [Kuhlman et al., 2013]. For example, one interesting proposal is that the increased inhibition enhances the visual-to-background activity ratio [Toyoizumi et al., 2013], while another theory proposed that increasing inhibition favoured more coherent inputs over stronger inputs [Kuhlman et al., 2010]. It is possible that multiple of these mechanisms play a role in OD plasticity. In our model, the background activity is crucial to model the MI since it allows us to increase the impact of MI on the neuronal firing rates. Moreover, assuming immature and adult levels of visual-to-background activity ratio lead to a better agreement of OD shifts between our model and experiments. Finally, adding more spontaneous activity in our model could counteract maturation if we assume that this spontaneous activity predominantly leads to synaptic depression, keeping the weights low and random. In this case maturation of V1 can only happen when the visual-to-spontaneous ratio is sufficiently high. This ratio could be gradually increased by the developmental changes in NMDA-receptor channels [Flint et al., 1997], nogo-receptors and myelination [McGee et al., 2005], inhibition [Toyoizumi et al., 2013] and changes in recurrent connectivity [Ko et al., 2013].

The recurrent excitatory connections in LII/III of our model are not critical for our results. These synapses allow us to reproduce the selectivity of excitatory connections, but the recurrence weak (see Methods). Therefore, with slightly different threshold values, similar results are obtained in a static networks without any E-to-E recurrence (Supp. Fig. 6c). It would be interesting to study the effect of richer recurrent dynamics [Rubin et al., 2015]. Furthermore, the I-to-E plasticity ensures that excitatory rates do not exceed a neuron-specific activity level. After deprivation, I-to-E connections first strengthen to counteract the depression of E-to-I connections and subsequently weaken again once the the inhibition recovers. In layer IV, a strengthening of I-to-E connections is observed after 2 days of MD [Maffei et al., 2006], however it is unclear whether these connections weaken again for longer deprivations.

The end of the critical period is much less understood. Experiments suggest that the adult levels of inhibition are reached during the CP [Kuhlman et al., 2011]. Furthermore, Kuhlman et al. [Kuhlman et al., 2013] showed that in adult mice no reduction of inhibition is observed after one day of MD. This readily leads to the assumption that the E-to-I plasticity, which is crucial in our model to observe OD plasticity, is somehow abolished. We therefore implemented the end of the critical period as a consolidation of the E-to-I plasticity, which could be mediated by changes in the extracellular matrix. Indeed, perineuronal nets (PNNs), have been shown to develop around PV+ inhibitory neurons at the end of the critical period [Pizzorusso, 2002], and could affect the plasticity of synapses onto these PV+ neurons [Wang and Fawcett, 2012]. Furthermore, degrading the PNNs in adult animals restored a window for OD plasticity [Pizzorusso, 2002] and this removal is related to reduced inhibition [Lensjø et al., 2017]. Another possibility is that the E-to-I plasticity rule itself prevents a reduction of inhibition in adults, for example due to changes in firing rates or correlations. Also the amount of silent synapses has been linked to the ability of juvenile-like plasticity [Huang et al., 2015]. The end of the critical period is characterized by the pruning of most of these silent synapses and a loss of PSD-95 in the adult leads to an increase in silent synapses as well as a recovery of OD plasticity. However, even though it was shown that the AMPA-to-NMDA ratio of excitatory synapses onto PV+ interneurons was similar in wild-type and PSD95-KO mice, it is not clear how a loss of PSD-95 affects E-to-I plasticity.

Our model allows for certain predictions that can be experimentally tested. Firstly, we hypothesize that neurons showing a substantial OD shift after MD need to have a sufficient difference between the lowest and the highest ODI of the input synapses (Fig. 2 and Supp. Fig. 2). It could be possible that the active synapses have a narrower distribution, but that reducing the inhibition uncovers a broader distribution of silent synapses. In this way, both mechanisms discussed previously - the presence of silent synapses [Huang et al., 2015] and the reduction of inhibition [Kuhlman et al., 2013] - could contribute to OD plasticity. Moreover, in animals with columnar organisations of OD we would only expect a broad distribution to neurons on the edges between columns. Our model would then predict that only these neurons show a fast OD shift. Secondly, we introduce a low threshold in our plasticity rule separating no-plasticity from plasticity. Such a low threshold has been observed experimentally [Artola et al., 1990] and was implemented in a modified version of the BCM-rule [Artola and Singer, 1993]. The low threshold allows us to reproduce the effects of MI [Frenkel and Bear, 2004], the dependence of OD shift on ODI [Mrsic-Flogel et al., 2007] and the counter-intuitively shifting cells under MD [Rose et al., 2016], while providing an explanation to why the latter tend to have lower firing rates. Thirdly, we assume that the E-to-I plasticity rule depends on the excitatory population activity. In our case, this was implemented using different target firing rates under normal vision and after deprivation. Such a dependence on population activity has been observed in hippocampal I-to-E plasticity [Hartman et al., 2006], and it would be interesting to investigate whether and how E-to-I plasticity implements similar mechanisms. Finally, our model predicts that increasing all excitatory firing rates during the monocular deprivation leads to a reduced OD shift.

To conclude, in this article we describe a theory of the development of cortical layer II/III. We implemented a simplified network with biologically plausible learning rules, which is able to reproduce multiple experimental results. With our model, we propose that:

- A transient reduction in inhibition in turn boosting the open-eye activity out of an LTD regime - is required to shift the population response towards the open eye.
- A substantial level of inhibition is necessary to observe OD plasticity, enabling a subsequent and sufficient reduction of the inhibition. The level of inhibition in immature networks is still too low and therefore not effective at amplifying excitation.
- Different ODI input distributions and firing rates can account for the variability in individual OD shifts. A low threshold separating no-plasticity from LTD is sufficient to account for counterintuitive shifts in neurons firing at lower rates.
- In adult animals, the reduction of inhibition is not observed [Kuhlman et al., 2013]. We hypothesize that network changes during the CP ultimately prevent E-to-I plasticity. These changes could for example be consolidation of E-to-I weights, changes in firing rates and/or changes in correlations.

## Methods

All LII/III neurons are modelled as rate units, where

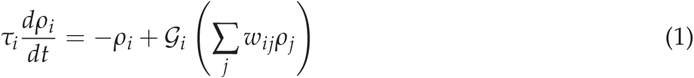

Here, *ρ_i_* denotes the firing rate of neuron *i*, *τ_i_* is the integration time constant and 𝒢_*i*_ is the gain function. The summation over index *j* runs over all presynaptic neurons, and we assume *wij* positive in case of excitatory presynaptic neurons and negative in case of inhibitory presynaptic neurons. In all our simulations, we used a linear gain function 𝒢(*x*) = *g* · *x* with slope *g* = 0.3 for all neurons. Finally, we used a time constant *τ_e_* = *τ*_*i*_ = 1ms.

### 1. Simplified Model

For our simplified model, we simulated 1250 presynaptic excitatory neurons, mimicking layer IV inputs, one postsynaptic inhibitory neuron and one postsynaptic excitatory neuron, mimicking layer II/III neurons. The inputs to the layer IV neurons were modelled as two visual inputs and a background input. The visual inputs represent both eyes and are modelled as step currents 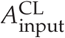 and 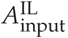, and the background input is an additional step current *B*. The visual input currents are multiplied by a weight in order to generate different ocular dominances. The weights were generated as follows:

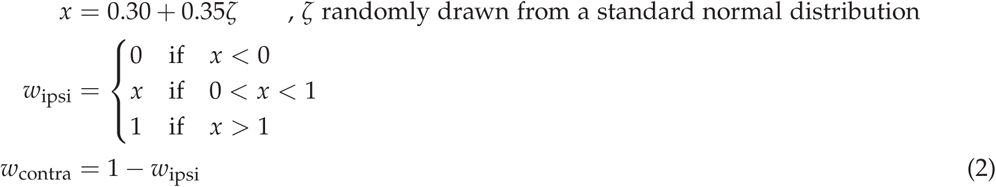

In other words, for each layer IV neuron, a random number was generated from a normal distribution with mean 0.30 and standard deviation 0.35. The resulting random numbers were rectified, i.e. negative numbers were set to 0, and numbers larger than 1 were set to 1. These numbers were chosen as the weights for the ipsilateral inputs to the layer IV neurons, while one minus the ipsilateral weights were the contralateral weights. In this way, each layer IV neuron received an equal amount of input, but with different ocular dominance. An example of the OD distribution for 1000 layer IV neurons is shown in Supp. Fig. 6a.

From the 1250 layer IV neurons, 250 randomly chosen connections are made to the layer II/III excitatory and inhibitory neurons. In this way, the layer II/III excitatory neurons receives inputs with a range of different ODIs. All inhibitory neurons received inputs from 150 most contralateral and 100 most ipsilateral neurons.

For simulations of Fig. 1 the feedforward E-to-E connections from layer IV to the layer II/III neuron (*w*_EE_) are plastic under the learning rule given by the following equation

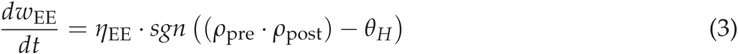

The parameter *θ_H_* is a threshold separating synaptic depression from synaptic potentiation. For simulations of Fig. 2, an extra threshold is introduced

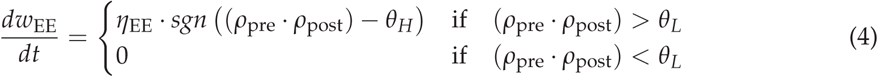

This learning rule is shown schematically in Fig. 3b. Here, the parameter *θ*_*H*_ is a threshold separating synaptic depression from synaptic potentiation, while *θ_L_* is a threshold below which no plasticity occurs. In both equations 3 and 4, *η*_EE_ is the learning rate and the function *sgn*(*x*) denotes the sign function and is equal to 1 if *x* > 0, equal to 0 if *x* = 0 and equal to -1 if *x* < 0. The plastic weights are constraint by hard lower and upper bounds, and all other weights (E-to-I, I-to-E) are static.

We start the simulations with the *w*_*EE*_ at the upper bound. The duration of the simulations is 10s, with timesteps of 1ms. We alternated activation of the inputs to the layer IV neurons (20ms at a constant value 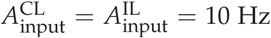 and a background activation *B* = 10Hz) with disactivation 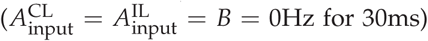. After 500ms, we simulated deprivation. In case of monocular deprivation, we reduced the contralateral input to zero, 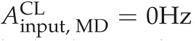. In the case of monocular inactivation, we reduced the contralateral input and the background input to zero, 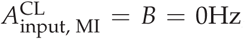, and in the case of binocular deprivation, we reduced the contra-and ipsilateral inputs to zero, 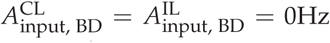. The values of all parameters used during the simulation can be found in Table 1.

**Table 1:** model parameters

|  |  |  |
| --- | --- | --- |
| Timestep | $dt$ | 1 ms |
| Gain function (slope) | $g_e = g_i$ | 0.3 |
| E time constant | $\tau_e$ | 1 ms |
| I time constant | $\tau_i$ | 1 ms |
| Nr. of L II/III exc. neurons | $NE_{L3}$ | 100 |
| Nr. of L IV exc. neurons | $NE_{L4}$ | 1000 |
| Nr. of L II/III inh. neurons | $NI_{L3}$ | 20 |
| Nr. of ffw connections per input group | $N_{ffw}$ | 50 |
| Max. E to E weight | $w_{EE}^{\max}$ | $1/N_{ffw}$ |
| Min. E to E weight | $w_{EE}^{\min}$ | $w_{EE}^{\max}/2000$ |
| High threshold EE | $\theta_H$ | $26 \text{ Hz}^2$ |
| Low threshold EE | $\theta_L$ | $15 \text{ Hz}^2$ |
| Learning rate EE | $\eta_{EE}$ | $12 \cdot 10^{-5} \text{ Hz}^{-2}$ |
| Max. I to E weight | $w_{EI}^{\max}$ | ∞ |
| Min. I to E weight | $w_{EI}^{\min}$ | $0.06 / NI_{L3}$ |
| High threshold I to E | $\phi_H$ | mean: 6 Hz (see methods) |
| Low threshold I to E | $\phi_L$ | $\phi_H - 0.75 \text{ Hz}$ |
| Learning rate I to E | $\eta_{EI}$ | $3.75 \cdot 10^{-4} \text{ Hz}^{-2}$ |
| Max. E to I weight | $w_{IE}^{\max}$ | $1/N_{ffw}$ |
| Min. E to I weight | $w_{IE}^{\min}$ | $w_{IE}^{\max}/12.5$ |
| Learning rate E to I | $\eta_{IE}$ | $3.75 \cdot 10^{-7} \text{ Hz}^{-3}$ |
| E to I target rate (normal vision) | $\rho_{\text{target}}$ | 6 Hz |
| BCM time constant | $\tau_{\text{avg}}$ | 12500 ms |
| Visual Inputs to L IV | $A_{\text{input}}^{\text{CL}} = A_{\text{input}}^{\text{IL}}$ | 10 Hz |
| Background input | $A_{\text{input}}^{\text{CL}} = A_{\text{input}}^{\text{IL}}$ | 10 Hz |

### Network

We simulated 1000 layer IV neurons, 100 layer II/III excitatory neurons and 20 layer II/III inhibitory neurons. The layer IV neurons are divided into five groups of 200, each representing a different input feature (e.g. a different orientation). Similarly, the layer II/III excitatory neurons are divided into five groups of 20 neurons. Each layer II/III neuron received 50 connections from each of the five layer IV groups (hence 250 feedforward synapses in total). However, connections from layer IV neurons of the same group were initialized at 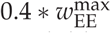, while other feedforward connections were initialized ten times smaller. In this way, an initial orientation preference at eye-opening was mimicked [Ko et al., 2013].

Moreover, the feedforward connections were not chosen randomly. If we would randomly pick these inputs, the probability to have very ipsilaterally or contralaterally dominated neurons in layer II/III is low, and instead all neurons would have an ODI close to the layer IV population ODI (Supp. Fig. 6b). Therefore, the connections were set to result in a broader ODI distribution in layer II/III. We outline below how we set 50 feedforward connections from one group of 200 layer IV neurons. The same is valid for the four other input groups representing the other orientations. The 200 layer IV neurons representing one orientation are divided into groups according to ocular dominance, *O*_1_ to *O*_5_. We then ensured a broad ODI distribution in layer II/III as follows. We choose *x* layer II/III neurons and randomly make *y_i_* connections from *O_i_* to these neurons, with *x* and *y_i_* given below.

| $x$ | $y_1$ | $y_2$ | $y_3$ | $y_4$ | $y_5$ |
| --- | --- | --- | --- | --- | --- |
| 10 | 5 | 5 | 5 | 10 | 25 |
| 10 | 5 | 10 | 10 | 10 | 15 |
| 20 | 5 | 10 | 10 | 10 | 10 |
| 20 | 10 | 15 | 10 | 10 | 5 |
| 20 | 10 | 15 | 15 | 5 | 5 |
| 20 | 25 | 10 | 5 | 5 | 5 |

This ensured both a broad distribution in layer II/III neurons (Fig. 4a), as well as enough contralaterally and ipsilaterally dominated inputs to each layer II/III neuron. The latter is important to be able to observe an OD shift in our model (see Results).

Recurrent E-to-E connections between the 100 layer II/III neurons were initialized randomly, as observed at eye-opening [Ko et al., 2013]. To this end, we picked random numbers from a normal distribution with mean and standard deviation equal to 5% of the maximum weight, and ensured values below the minimum weight were reset at this minimum weight.

For each of the 20 inhibitory neurons we randomly picked one of the input groups representing an orientation, to which it made 50 strong initial connections equal to the maximum weight, while 50 connections from each of the other input groups were initialized at the minimum weight. Similarly to the simplified model, all inhibitory neurons received inputs from 30 most contralateral and 20 most ipsilateral neurons in each input group.

Finally, all feedforward input strengths from layer IV to layer II/III excitatory and inhibitory neurons are multiplied by a factor of 5 to mimic a larger input population.

The feedforward and recurrent E-to-E plasticity rule is given by equation 4. This learning rule is shown schematically in Fig. 3b. The thresholds are chosen such that the open eye inputs after deprivation are above the highest threshold, while the closed eye inputs are under this threshold. Furthermore, we choose the low threshold such that all inputs are below this threshold after monocular inactivation. Finally, we show that our results are robust against the exact choice of thresholds (Supp. Fig. 5a).

The E-to-I plasticity rule is a BCM-type rule [Bienenstock et al., 1982], given by

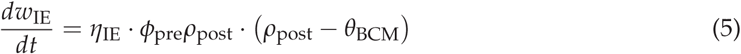

Following the results from [Kuhlman et al., 2013], the rule should ensure a quick depression of E-to-I inputs from both eyes and a subsequent recovery of mainly the open eye. Therefore, we choose no dependence on presynaptic rate for depression, *ϕ*_pre_ = 2Hz, while we choose a dependence on presynaptic rate for potentiation, *ϕ*_pre_ = *ρ*_pre_ if *ρ*_pre_ > 3.2Hz and zero otherwise.

Finally, to ensure the inhibition reaches roughly half of its value after deprivation as observed in [Kuhlman et al., 2013], we raise the minimum E-to-I weight to 0.4 of the maximum value. This E-to-I rule is such as to reproduce experimentally observed inhibitory activity (Fig. 4c), but we do not have enough data to further constrain the rule.

*θ*_BCM_ is a sliding threshold given by (< *ρ*_post_ >)^2^/*ρ*_target_. The average of the peak postsynaptic firing rate < *ρ*_post_ > is calculated online by low-pass filtering with a long time constant,

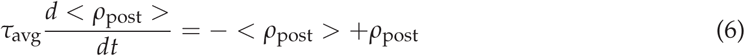

The *ρ*_target_ is a target firing rate for the inhibitory neurons, and was chosen to be 6 [a.u.] for normal vision, but reduced to 10% of this value after deprivation. Finally, the I-to-E plasticity rule is based on the rule in Vogels et al. [Vogels et al., 2011].

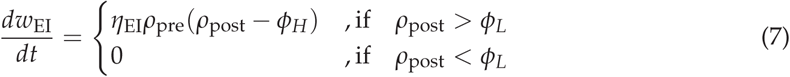

By choosing the lower threshold for plasticity *ϕ_L_* at a high value, close to but below *ϕ_H_*, this rule limits the maximum firing rates of the excitatory neurons but does not homeostatically increase the firing rate when sudden drops occur (for example after MD).

To simulate heterogeneous firing rates, for each layer II/III neuron we generate a random number *x* = 1 + 0.1*ζ*, with *ζ* drawn from a standard normal distribution. We then multiply all the rates of the presynaptic neurons of a layer II/III neuron by the respective random number *x* (thus either increasing or decreasing all presynaptic rates), while also multiplying the *ϕ_H_* of equation 7 for all I-to-E synapses to this layer II/III neuron with *x* (thus increasing or decreasing the maximal postsynaptic rate by an equal amount as the presynaptic rates). The threshold *ϕ_L_* was always a fixed amount lower than *ϕ_H_*.

The first phase of the simulation lasts 50s. In this phase, the excitatory and inhibitory connections develop and reach either the maximum or minimum bound. Furthermore, MD does not lead to OD shifts because initially, the inhibition is weak and reducing inhibition cannot enhance the excitation sufficiently.

After this first phase ensured stationary weights and a sufficiently high sliding threshold, we simulate the deprivation. This second phase lasts 50s. Similarly as in the simplified model, we model MD by reducing the closed-eye input to layer IV to zero, BD by reducing both ipsilateral and contralateral inputs to zero, and MI by setting the contralateral input and the background input to zero. To simulate adult animals, we do not allow any plasticity from E-to-I connections. To simulate pre-CP deprivation, we reduce the first phase of the simulation to 2s instead of 50s, ensuring that the excitatory and inhibitory weights are still low. Furthermore, we simulate the developmental shift in visual inputs strength by (1) increasing the background input from 10 to 15, and reducing the visual inputs from 10 to 5 in the pre-CP case, (2) decreasing the background input from 10 to 5 and increasing the visual inputs from 10 to 15 in the adult case.

### 3. ODI and selectivity index

The ocular dominance index (ODI) was calculated as

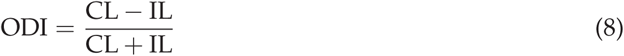

where CL stands for the maximum response to a contralateral visual input, and IL stands for the maximum response to an ipsilateral visual input. In this way, a neuron with ODI=1 is completely monocular for the contralateral eye, while a neuron with ODI=-1 is completely monocular for the ipsilateral eye.

The input selectivity index (SI) was calculated as one minus the circular variance. We first calculated the maximal response *a_j_* of a neuron to each of the *N*=5 inputs, and sorted these responses from large to small (*a*_1_ is the largest and *a*_5_ the smallest). Then, we calculated

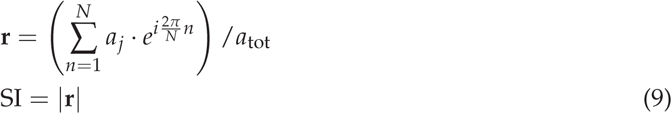

where 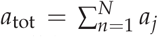. In this way, a neuron that is active for one input but silent for all other inputs will have an SI equal to 1, while an input that is equally active for all inputs will have an SI equal to 0.

The code for our models and all simulations herein will be available on ModelDB after publication (https://senselab.med.yale.edu/modeldb/).

## Acknowledgements

This work has been supported by the Wellcome Trust and the BBSRC.

## Supplementary Information

**Supplementary Figure 1.**
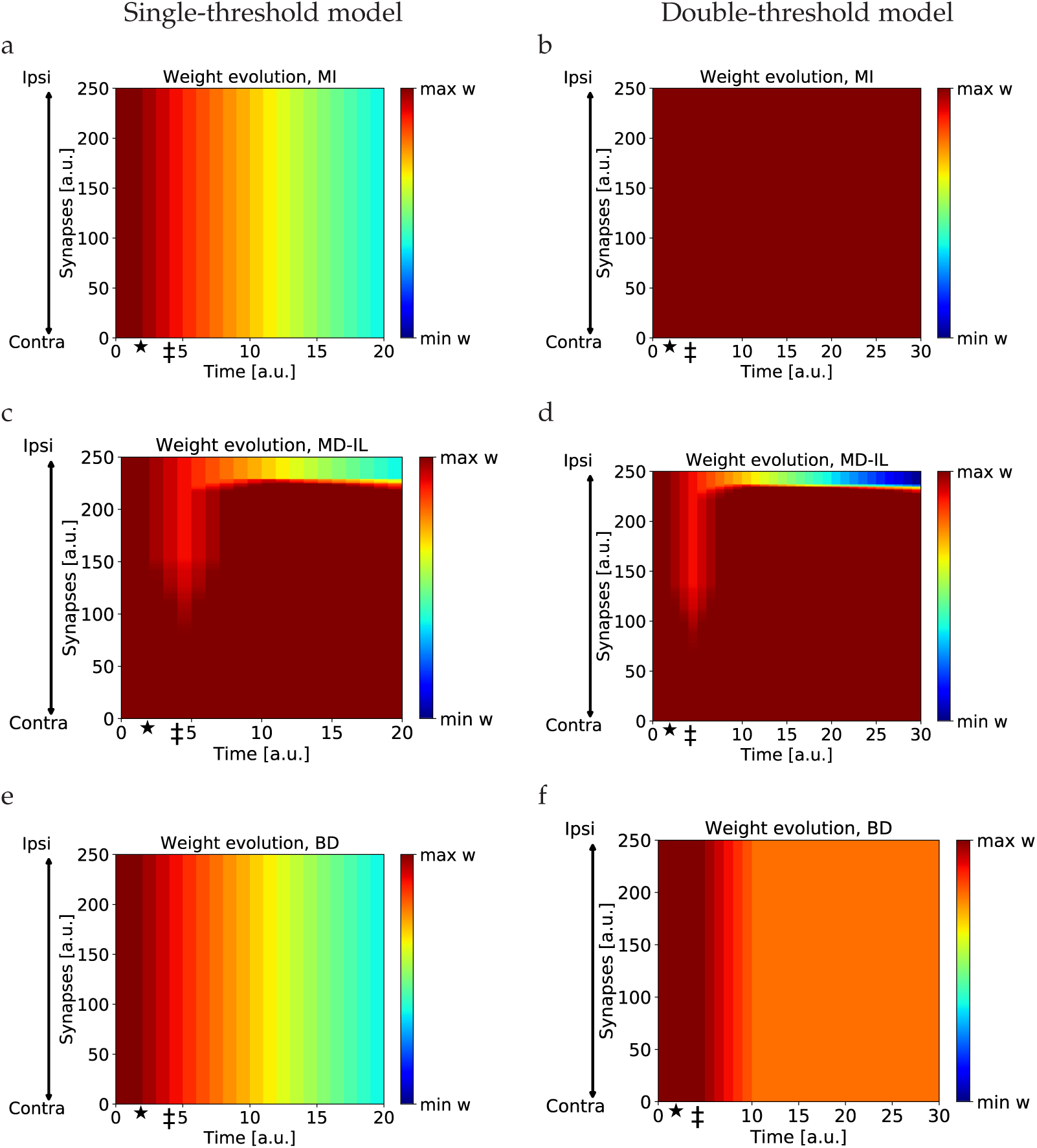
Evolution of weights. (a), (c), (e) Evolution of synaptic weights over time for the single-threshold model, in the cases of MI, MD-IL and MD-BD respectively. The star denotes the onset of deprivation, the double dagger denotes the onset of reduced inhibition. (b), (d), (f) Evolution of synaptic weights over time for the double-threshold model, in the cases of MI, MD-IL and BD. The star denotes the onset of deprivation, the double dagger denotes the onset of reduced inhibition.

**Supplementary Figure 2.**
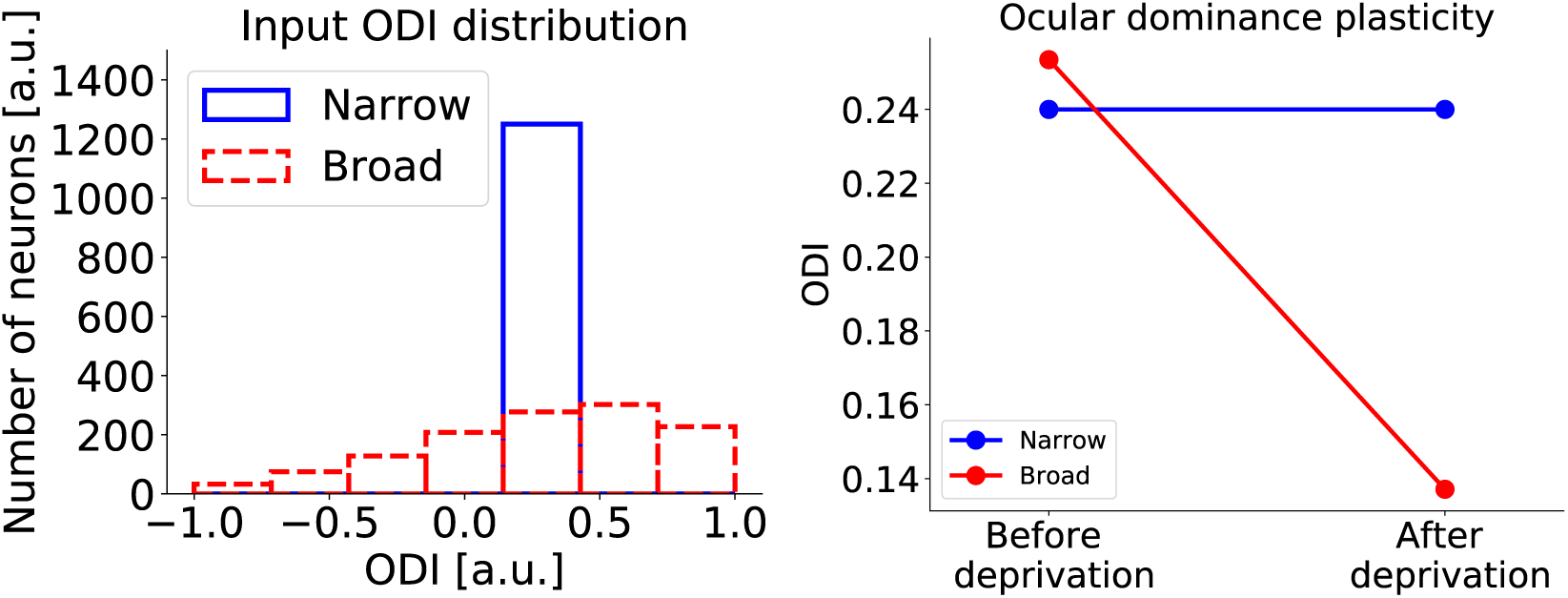
How the input distribution affects OD shifts. (a) A layer II/III neuron receives inputs with either a narrow ODI distribution (blue) or a broad distribution (red). (b) Layer II/III neurons with similar ODI index only show an OD shift after MD when the inputs have a broad ODI distribution.

**Supplementary Figure 3.**
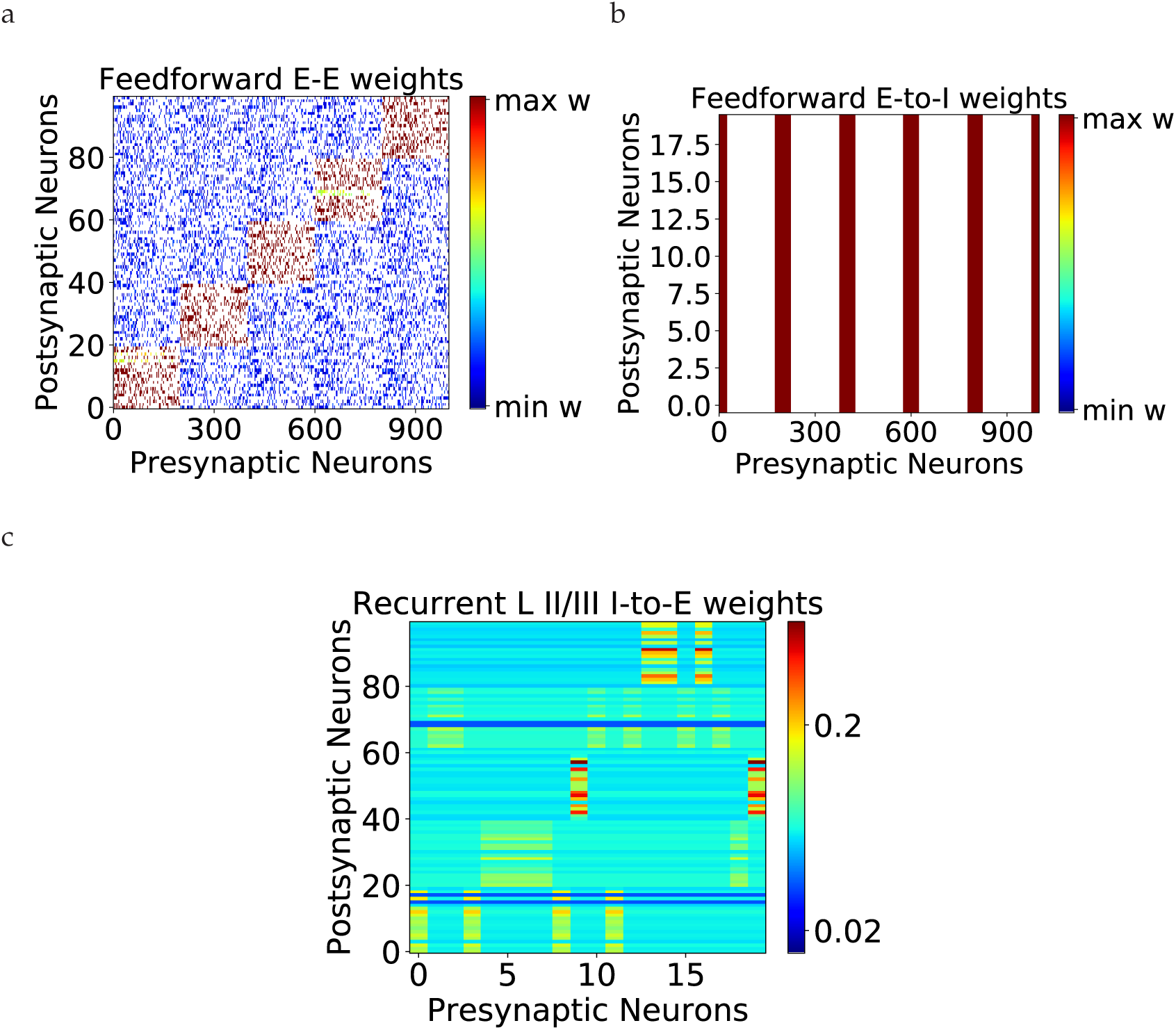
Synaptic Weights after first learning phase. (a) Feedforward E-to-E weights are specific, only synapses from one input group are strong, while other feedforward inputs are weak. Only 50 feedforward connections per input group are made, white denotes no connection (see Methods).(B) Feedforward E-to-I weights are unspecific, synapses from all input groups (200 neurons) are at the maximum bound. Only 50 feedforward connections per input group are made, white denotes no connection (see Methods). (c) Recurrent I-to-E weights after the first learning phase.

**Supplementary Figure 4.**
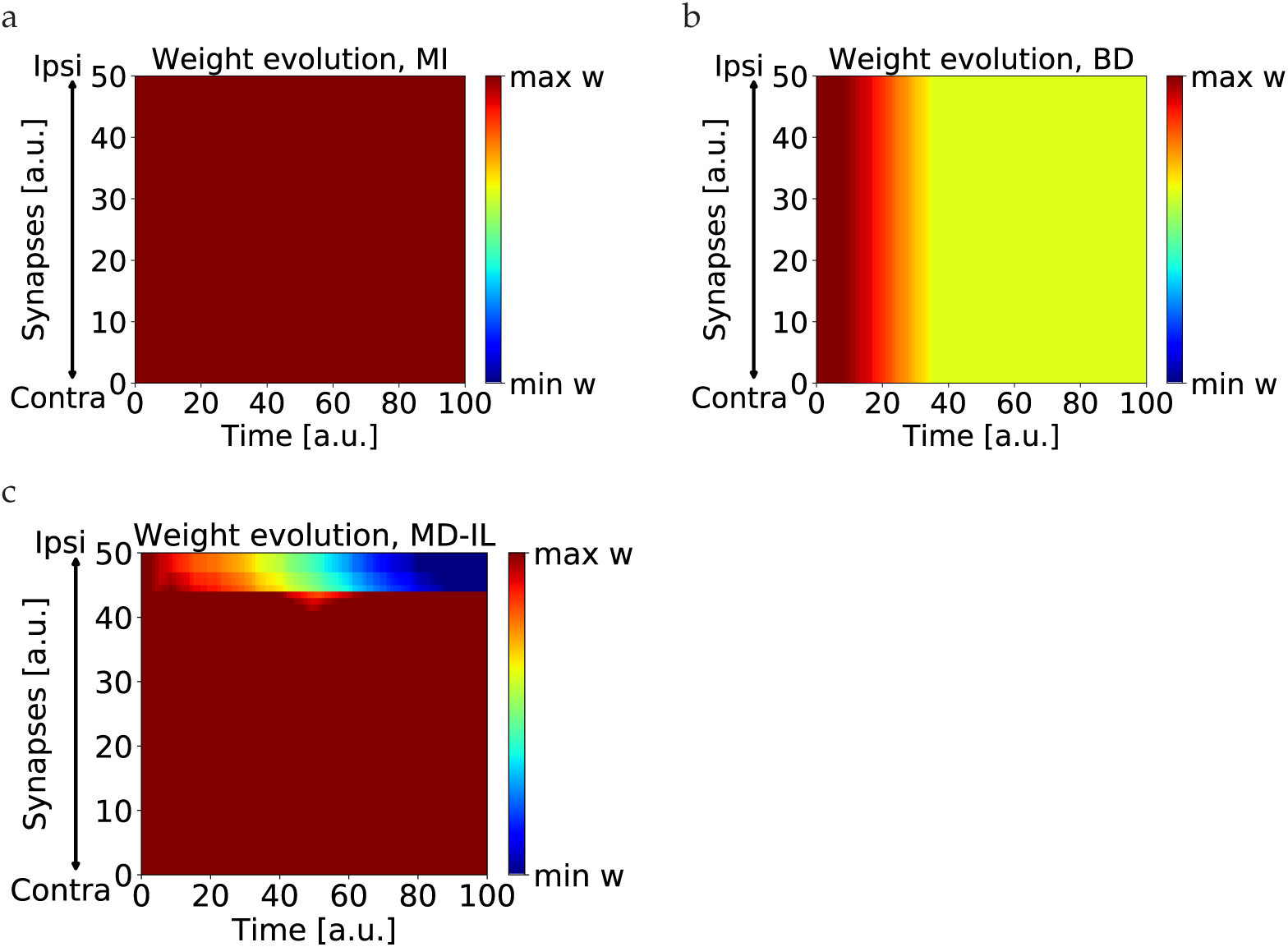
Evolution of network weights. (a), (b), (c) Evolution of synaptic weights over time for one neuron in the network, in the cases of MI, BD, MD-IL.

**Supplementary Figure 5.**
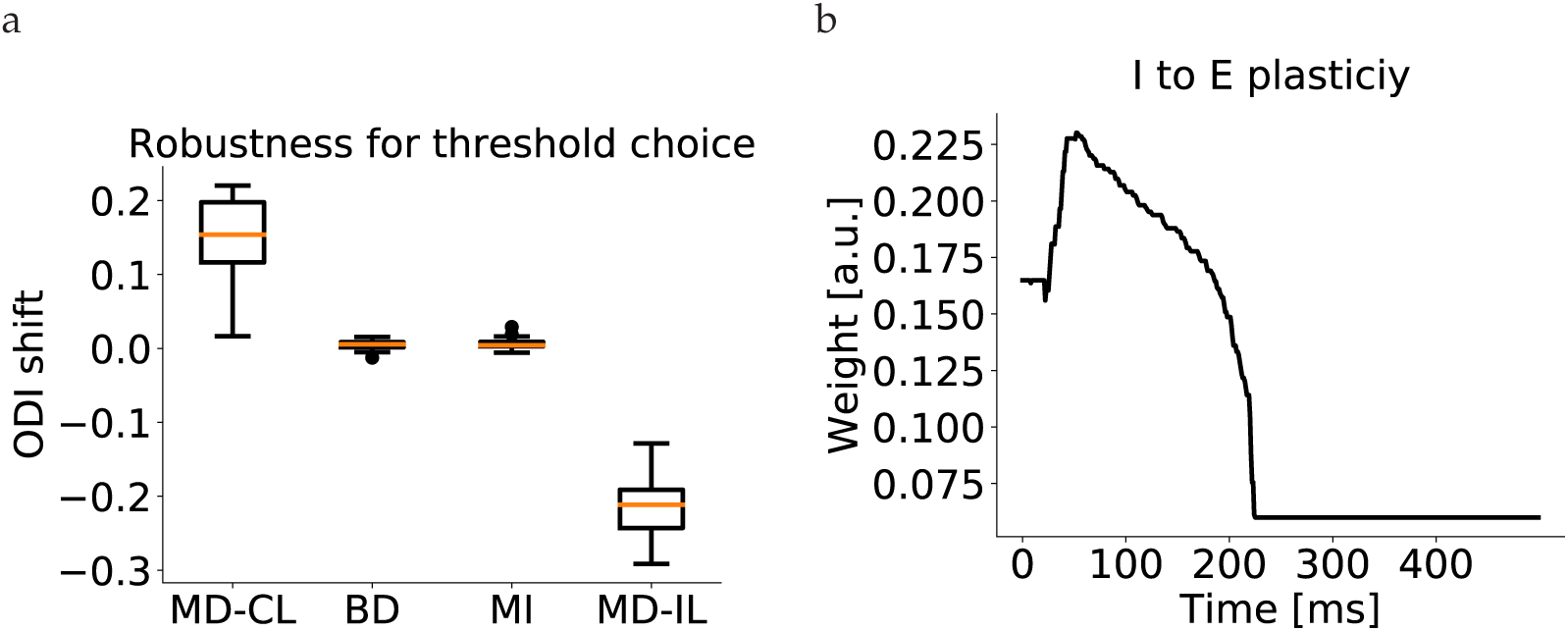
Thresholds and I-to-E weights. (a) Our results are robust for changes in threshold values. The boxplots show the distribution of population ODI shifts after 50 simulations for all types of deprivation. In each simulation, a random normal value for *θ_H_* and *θ_L_* is chosen with as mean the usual values (see Methods) and standard deviation 10% of these values. (b) Evolution of the I-to-E weights after deprivation. Potentiation is observed to partially counteract the reduction of inhibition in E-to-I connections, followed by depression to a minimum bound when the inhibition recovers.

**Supplementary Figure 6.**
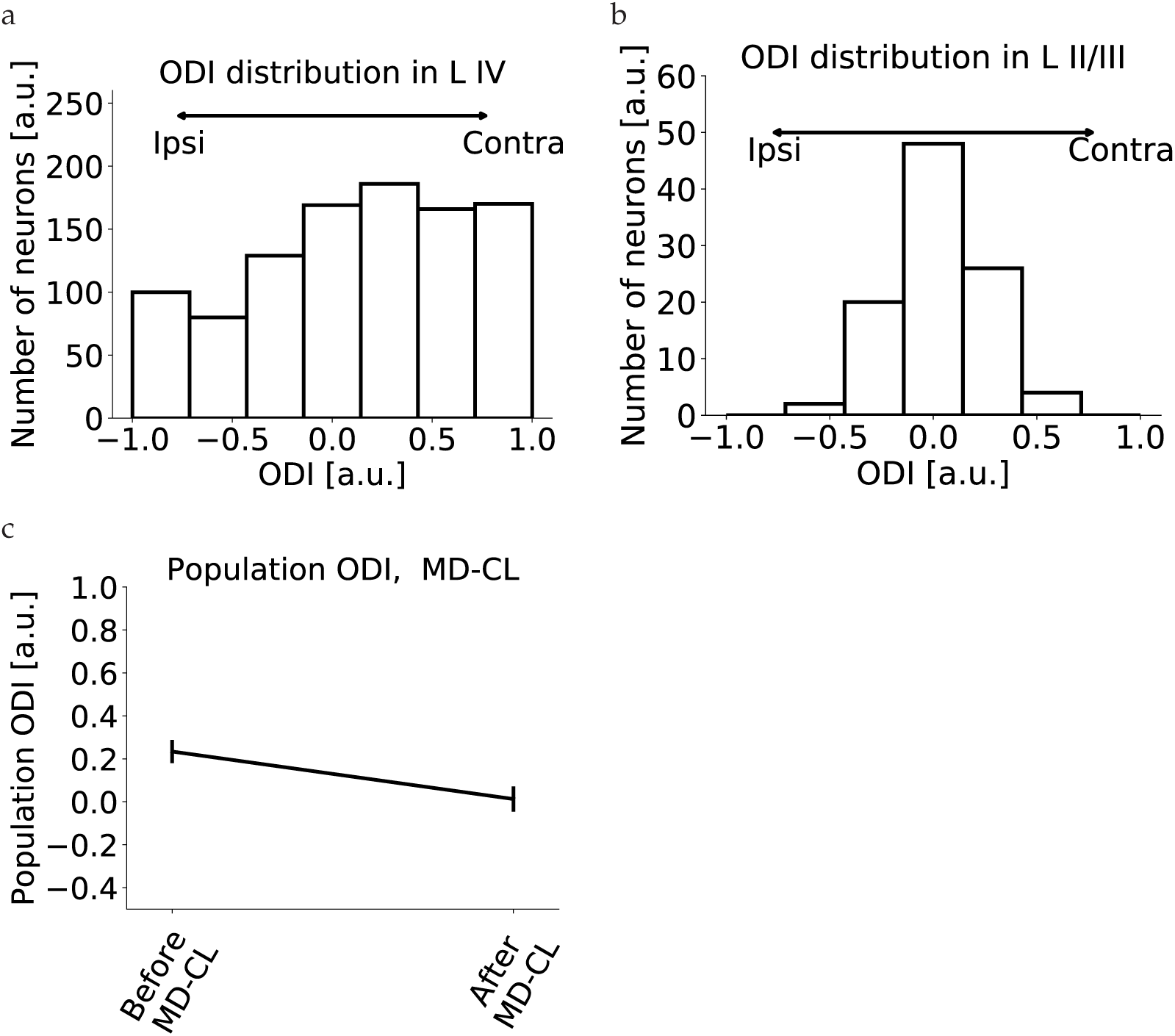
L IV and LII/II recurrence. (a) Example of ODI distribution in layer IV. (b) Randomly sampling 50 connections from the layer IV population leads to a narrow distribution in layer II/III. (C) Recurrent layer II/III connections are not crucial in our model. Similar results are obtained by reducing the *θ_H_* and the *θ_L_* by 1.

